## Supplemental Figures 1 and 2 for "Limited evidence of a genetic basis for sex determination in the common creek chub, *Semotilus atromaculatus*"

### 1 Supplement

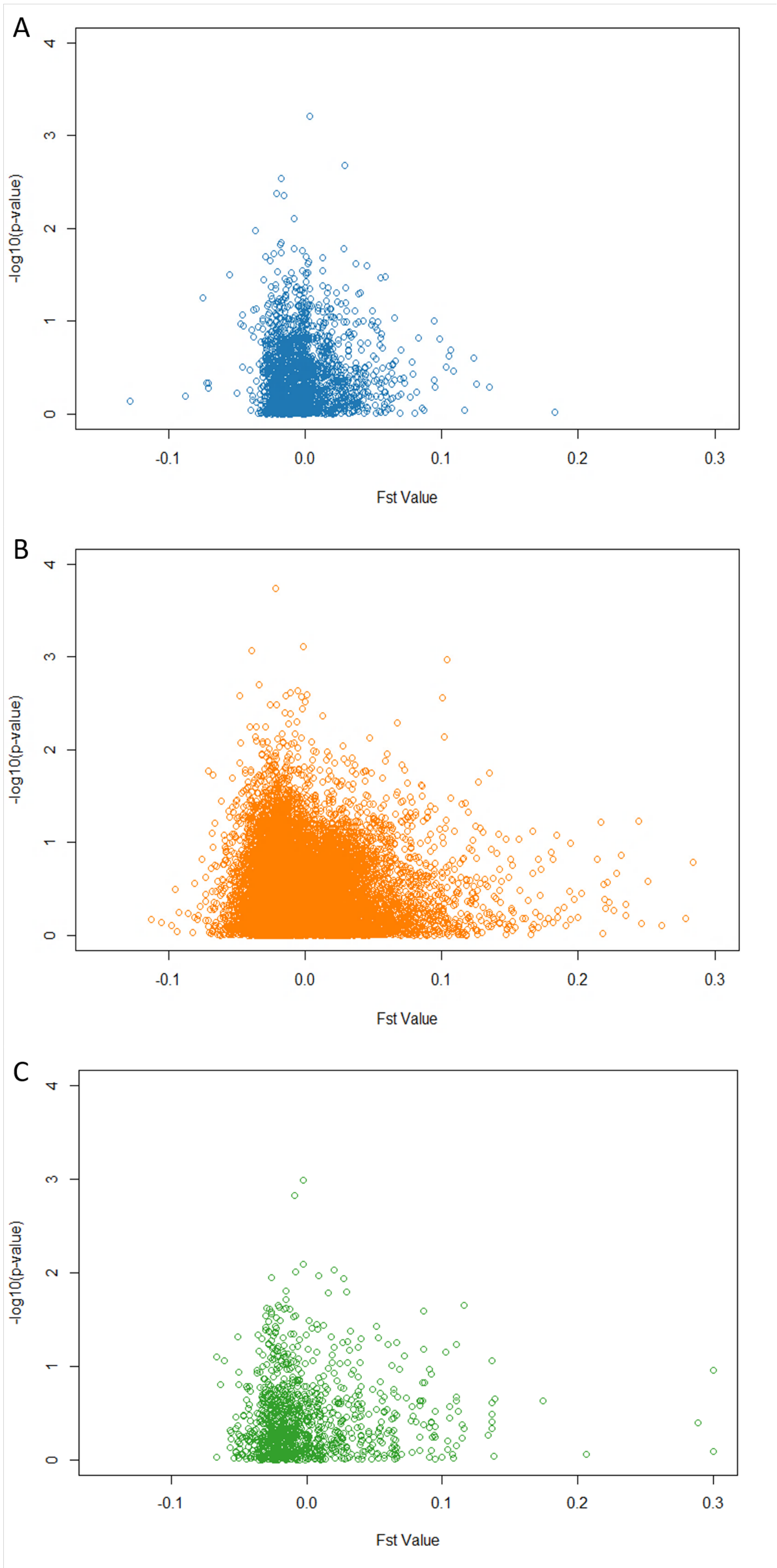


Figure S1: Scatter plot of association of markers from FST analysis and GWAS. FST value is displayed on the x-axis and the -log10(p-value) for each marker's association with sex is displayed on the y-axis. A: 3052-marker data set. B: 16,020-marker data set. C: 1132-marker data set.

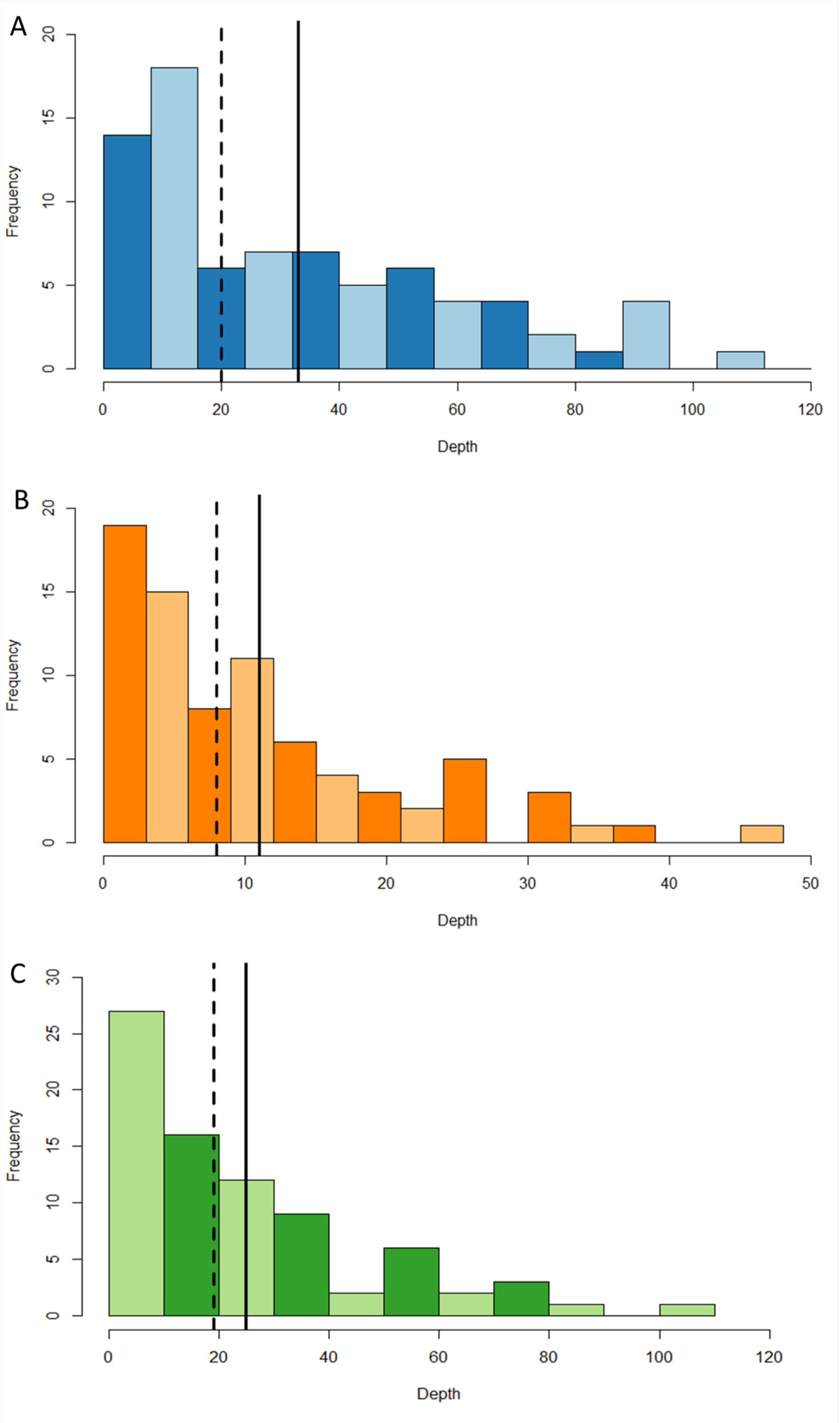


Figure S2: Histogram of mean read depth per marker, for all 79 individuals. Mean read depth is displayed on the x-axis and number of individuals is displayed on the y-axis. Solid vertical line denotes mean read depth and dashed vertical line denotes median read depth. A: 3052-marker data set. B: 16,020-marker data set. C: 1132-marker data set.
