## Supplement .tex file for "Limited evidence of a genetic basis for sex determination in the common creek chub, *Semotilus atromaculatus*": main.tex

\documentclass[letterpaper, 12pt]{article}

%---------------------------------------- + ---- + ---- + ---- + ---- + ----
\setlength{\parskip}{1ex plus0.5ex minus0.5ex}
\setlength{\textheight}{9.0in}
\setlength{\textwidth}{6.5in}
\setlength{\topmargin}{-0.5in} %% remove if using latex, rather than pdflatex
\setlength{\oddsidemargin}{0in}
%---------------------------------------- + ---- + ---- + ---- + ---- + ----

\newlength{\dblfigwidth}
\setlength{\dblfigwidth}{\textwidth}

\newlength{\figwidth}
\setlength{\figwidth}{4in}

\setlength{\fboxsep}{3mm}
%to include figures
\usepackage{graphicx}
%to get author-year citations
\usepackage{natbib}
\bibpunct{(}{)}{,}{}{}{,}

%% to include parenthetical URLs
\usepackage{url}
%% to include tables that are >1 page
\usepackage{longtable}
% for line numbering
\usepackage{lineno}
%\usepackage{times}
\usepackage{setspace}
%\doublespacing
\usepackage{rotating}
% ---------------
% to convert from latex to html:
% htlatex filename

\usepackage{cancel}
%strike through text

% to get page-numbering in upper-right corner
\pagestyle{myheadings}

\begin{document}
\begin{linenumbers}

%-----------------------------------------------------------------------------------

\section{Supplement}
\setcounter{table}{0} \renewcommand{\thetable}{S\arabic{table}}
\setcounter{figure}{0} \renewcommand{\thefigure}{S\arabic{figure}}

\clearpage
%-----------------------------------------------------------------------------------

\begin{figure}
    \centering
    \includegraphics[scale=0.5, angle=0]{figures/gwas_fst_vertical.png}
    \caption{Scatter plot of association of markers from FST analysis and GWAS. FST value is displayed on the x-axis and the -log10(p-value) for each marker’s association with sex is displayed on the y-axis. A: 3052-marker data set. B: 16,020-marker data set.  C: 1132-marker data set.}
    \label{fig:gwas_fst} 
\end{figure}


\clearpage
%-----------------------------------------------------------------------------------


\begin{figure}
    \centering
    \includegraphics[scale=0.5, angle=0]{figures/vcf_depth_vertical.png}
    \caption{Histogram of mean read depth per marker, for all 79 individuals. Mean read depth is displayed on the x-axis and number of individuals is displayed on the y-axis. Solid vertical line denotes mean read depth and dashed vertical line denotes median read depth.  A: 3052-marker data set. B: 16,020-marker data set. C: 1132-marker data set.}
    \label{fig:vcf} 
\end{figure}


\clearpage

%-----------------------------------------------------------------------------------

\end{linenumbers}
\end{document}
