## Supplementary material for "Limited evidence of a genetic basis for sex determination in the common creek chub, *Semotilus atromaculatus*": Manuscript .tex file: main.tex

\documentclass[letterpaper, 12pt]{article}

%---------------------------------------- + ---- + ---- + ---- + ---- + ----
\setlength{\parskip}{1ex plus0.5ex minus0.5ex}
\setlength{\textheight}{9.0in}
\setlength{\textwidth}{6.5in}
\setlength{\topmargin}{-0.5in} %% remove if using latex, rather than pdflatex
\setlength{\oddsidemargin}{0in}
%---------------------------------------- + ---- + ---- + ---- + ---- + ----

\newlength{\dblfigwidth}
\setlength{\dblfigwidth}{\textwidth}

\newlength{\figwidth}
\setlength{\figwidth}{4in}

\setlength{\fboxsep}{3mm}
%to include figures
\usepackage{graphicx}
%to get author-year citations
\usepackage{natbib}
\bibpunct{(}{)}{,}{}{}{,}

%% to include parenthetical URLs
\usepackage{url}
%% to include tables that are >1 page
\usepackage{longtable}
% for line numbering
\usepackage{lineno}
%\usepackage{times}
\usepackage{setspace}
%\doublespacing
\usepackage{rotating}
% ---------------
% to convert from latex to html:
% htlatex filename

\usepackage{cancel}
%strike through text

% to get page-numbering in upper-right corner
\pagestyle{myheadings}

\begin{document}

\noindent{\large {\bf Limited evidence of a genetic basis for sex determination in the common creek chub, \emph{Semotilus atromaculatus}}}

\medskip


\noindent {\large Amanda V. Meuser$^{1,2}$, Cassandre B. Pyne$^{2}$, Elizabeth G. Mandeville$^{2}$}


\bigskip


\noindent $^1$ \parbox[t]{6.5in}{Department of Molecular and Cellular Biology, University of Guelph, Guelph, Ontario, Canada}
  
\noindent $^2$ \parbox[t]{6.5in}{Department of Integrative Biology, University of Guelph, Guelph, Ontario, Canada}


\vspace{1in}

\noindent \textbf{Corresponding author}: \parbox[t]{4in}{Amanda Meuser\\
  University of Guelph\\
  50 Stone Road East\\
  Guelph, Ontario N1G 2W1, Canada\\
 \\
}

\bigskip

\noindent \textbf{ORCID IDs:} Amanda V. Meuser - 0000-0001-9554-2002, Cassandre B. Pyne - 0000-0002-2344-2800, Elizabeth G. Mandeville - 0000-0002-9461-9140


\medskip

\noindent \textbf{Running title}: Sex determination in creek chub

\medskip

\noindent \textbf{Keywords}: Sex determination, sex chromosomes, creek chub, \emph{Semotilus atromaculatus}, genotyping-by-sequencing, RADSex, GEMMA, F$_{ST}$, linkage disequilibrium

\medskip

\noindent \textbf{Author contributions}:
AVM and EGM planned research, designed analysis strategy, and implemented bioinformatics workflows. CBP contributed to analyses. All authors wrote and revised the manuscript. 

\medskip

\noindent \textbf{Conflict of Interest}: The authors have no conflict of interest to declare. 

\medskip

\noindent \textbf{Acknowledgements}: 
Thanks to the McCann Lab at the University of Guelph for collection of the Creek Chub samples. Permission to collect was also granted from private landowners and the following conservation areas: Hawkins Trail Conservation Area, Kettle Creek Conservation Area, Watson Conservation Area, Backus Woods Conservation Area. 
This research was undertaken thanks in part to funding from the Canada First Research Excellence Fund Food From Thought program at the University of Guelph. 
Computing was accomplished through a RRG Allocation on Compute Canada’s Graham cluster to E.G. Mandeville. 
We thank Daniel Fabrizio for assistance with DNA extractions. This manuscript was improved by comments from Eryn McFarlane, Ben Schultz, Jillian Campbell, Amy Pitura, Eric Sartor, and Erza Dafota.

\medskip


\begin{linenumbers}

%-----------------------------------------------------------------------------------

\pagebreak
\noindent{\Large {\bf Limited evidence of a genetic basis for sex determination in the common creek chub, \emph{Semotilus atromaculatus}}}

\medskip

\section{Abstract}

%-----------------------------------------------------------------------------------


Sexual reproduction is almost universal in the animal kingdom; therefore, each species must have a mechanism for designating sex as male or female. Fish especially have a wide range of sex determining systems. While incompatible systems are thought to increase reproductive isolation, interspecific hybridization is common among groups such as cyprinid minnows, thus, studies such as this can provide insight into hybridization and evolutionary diversification of this clade. In the present study, we aimed to identify a genetic basis for sex determination in the common creek chub (\textit{Semotilus atromaculatus}) using genotyping-by-sequencing (GBS) data. No sex-associated markers were found by RADSex or a GWAS using GEMMA, however, our Weir and Cockerham locus-specific F$_{ST}$ analysis and discriminant analysis of principal components revealed some genetic differentiation between the sexes at several loci. While no explicit sex determination mechanism has been yet discovered in creek chub, these loci are potential candidates for future studies. This study also highlights technical challenges involved in studying sex determination in species with extremely variable mechanisms.


%-----------------------------------------------------------------------------------

\section{Introduction}

%-----------------------------------------------------------------------------------

Incompatible sex determination mechanisms or loci may lead to diversification of species and an increase in biodiversity, as it contributes to reproductive isolation. Conversely, genetically distinct species with compatible sex determination mechanisms may be able to hybridize, which could reduce overall biodiversity if either parent species is lost to genetic or demographic swamping \citep{Dowling1993, ONeill2018a, Todesco2016}. Understanding how sex determination plays a role in preventing or allowing species hybridization can shed light on the evolution of species-rich groups such as cyprinid minnows, as well as how anthropogenic disturbance may influence this evolution through hybridization in the future, as previously isolated species are being newly introduced into sympatry \citep{Grabenstein2018, Todesco2016}.

Sex determination mechanisms are extremely diverse, but can be broadly bisected into genetic sex determination (GSD) and environmental sex determination (ESD), for gonochoristic species (those with separate sexes) \citep{Bachtrog2014a, Charnov1977, Herpin2015, Heule2014}. In some cases of GSD, multiple genes or genetic regions across two or more chromosomes work in a threshold-dependent manner to determine sex. These are known as either polygenic sex determination or multiple sex chromosome systems, which may arise from chromosomal fissions and fusions \citep{Bachtrog2014a, Bulmer1982ModelsControl, Charnov1977, Faggion2019a, Heule2014, Kitano2012b, Kosswig1964, Vandeputte2007ALabrax}.

More commonly, however, GSD is under the control of a single locus, the master sex determining (MSD) gene \citep{Herpin2015, Heule2014, Pan2019, Pennell2018a}. An obvious example of GSD with an MSD gene is the heteromorphic XY sex chromosomes in mammals, universal across the clade. The male Y chromosome contains the gene Sry, which acts as a transcription factor for genes required to form the male phenotype \citep{Bachtrog2014a, Herpin2015, Kashimada2010}. Due to the lack of homology with the X chromosome, this area became isolated from recombination. Over time, mutations and gene loss have eroded the Y chromosome, such that it became cytologically distinct from its X counterpart \citep{Charlesworth1996TheCompensation, Charlesworth2017, Furman2020, Heule2014, Pennell2018a}. 

In contrast to the example of heteromorphic sex chromosomes, species with homomorphic sex chromosomes have the same sex determination genes in both males and females and instead possess different alleles which trigger genes for the development of distinct phenotypes \citep{Furman2020, Heule2014}. These sex chromosomes share a greater percentage of homology, are able to recombine to a greater extent, are subject to lower rates of degradation through gene loss, and remain cytologically indistinguishable from their counterpart \citep{Charlesworth1996TheCompensation, Heule2014, Pan2019}. Rates of linkage disequilibrium (LD), the measure of non-random assortment of alleles, can therefore be used to support evidence for potential sex determination regions, as markers present in these non-recombining areas would be expected to have high values of LD \citep{Charlesworth1996TheCompensation, Kamiya2012b, Pan2019}. For example, tiger pufferfish (\emph{Takifugu rubripes}) possess homomorphic sex chromosomes with alleles that differ by only a single nucleotide and recombine freely throughout this region of the genome \citep{Kamiya2012b}. Thus, homomorphic sex chromosomes are often more arduous to detect than heteromorphic, especially in fish, which vary greatly in sex determination systems from species to species \citep{Bachtrog2014a, Natri2019a, Pan2019}. 


In this study, we used data from individuals of known sex to investigate a potential genetic basis of sex determination in \emph{Semotilus atromaculatus} (the common creek chub), a cyprinid fish. While hermaphroditism is ten times more common in fish than other animal clades, gonochorism has been exclusively observed in creek chub, and most cyprinid fish are believed to have genetically determined sex \citep{Bachtrog2014a, Pennell2018a, Copes1978a}. Thus, only the mechanisms of sex determination for gonochoristic species were investigated. Several complementary genomic analyses were used on genotyping-by-sequencing (GBS) data, with the objective of determining areas of the genome that are associated with sex in creek chub. 


%-----------------------------------------------------------------------------------

\section{Materials and methods}

%-----------------------------------------------------------------------------------


\subsection{Experimental model and subject details}

Creek chub fish were collected from 14 agricultural streams across Southern Ontario in the summer of 2019. Fish were euthanized using an overdose of MS-222, then fin tissue was sampled and preserved in 95\% ethanol. 79 individuals were used in this study; 38 females and 41 males. Sex was determined through dissection. 

Sampling was done under the consent of Animal Use Protocol \#3682, approved by the University of Guelph Animal Care Committee (F. Laberge PI) and Ministry of Natural Resources Licence to Collect Fish for Scientific Purposes (K. McCann PI).

\subsection{DNA isolation and genomic sequencing}


We extracted DNA from each of the samples using the QIAGEN DNeasy Blood \& Tissue Kit (Qiagen, Inc.), according to the manufacturer's instructions. Following DNA extraction, we prepared genomic libraries using a reduced-representation genotyping by sequencing method \citep{Parchman2012} to prepare samples for high-throughput sequencing. DNA was digested with EcoRI and MseI restriction enzymes, then adaptors containing the adaptor sequence, barcode, cut site, and a protector base were ligated onto the fragments, before being amplified with Illumina PCR primers added onto either end \citep{Parchman2012}. After amplification, the DNA samples were sent to the Genomic Sequencing and Analysis Facility at the University of Texas at Austin. There, the samples were size selected for fragments 300-400bp in length using the Sage Science Blue Pippin, then sequenced with an Illumina NovaSeq 6000. All computing was accomplished via an allocation on Compute Canada’s high-performance computing cluster, Graham, with the exception of data visualization in RStudio on a local computer.


\subsection{Primary investigation of all GSD types}


We used several complementary methods to investigate the genetic basis of sex determination in creek chub. We first demultiplexed the raw FASTQ file using sabre (https://github.com/na
joshi/sabre), resulting in one FASTQ file for each individual fish. We performed no additional quality control on the reads. While there may be some unintended additional sequences remaining in the data (e.g. primers, adaptors), it is not expected to bias our results. Initial analysis of sex-associated markers in the creek chub genome was performed using RADSex, a computational workflow intended as an “all-in-one” GSD detection program and visualizer, with the assistance of the RStudio package {\tt sgtr} \citep{Feron2021a}. RADSex treats all markers as unassociated genotypes, rather than alleles of the marker, and searches for markers that are found in one sex exclusively, at a statistically significant level \citep{Feron2021a}. Thus, it is theoretically able to detect heteromorphic or homomorphic sex chromosomes, or polygenic sex determination loci, using only one workflow. Additionally, it does not require a genome for alignment, making it ideal for use with non-model organisms, and does not require imputation before use as it allows for missing data.

RADSex takes an input of a FASTQ file of demultiplexed reads for each individual and a file with a list of each individual and their sex. First, the software creates a table containing each marker and its read depth, then the table is filtered such that only markers with a certain read depth are retained \citep{Feron2021a}. To fit the low-coverage GBS data used for the present study, we set the depth threshold to one read. Next, RADSex identifies makers that are significantly associated with sex, using Pearson’s chi-squared test of independence \citep{Feron2021a}. 
We visualized the output in RStudio using the {\tt radsex\_distrib} function from the accompanying R package, {\tt sgtr}, which creates a tile plot (Fig.\ \ref{fig:radsex}).

\subsection{Identification of variant sites and filtering of VCF files}


As there is not yet a reference genome for creek chub, we constructed an artificial reference genome with CD-HIT \citep[as described in][]{LaCava2019}. We aligned FASTQ files to this reference and sorted into contigs using the Burrows-Wheeler Alignment tool (BWA) \citep[version 0.7.17]{Li2009b}. We identified variable genetic loci using {\tt bcftools mpileup} \citep[version 1.11]{Li2009b} and converted the resulting BCF  to a VCF file using SAMtools \citep[version 1.12]{Li2009c}. We then filtered the resulting VCF using VCFtools \citep{Danecek2011TheVCFtools}. We produced two different filtered VCFs to fit our analyses.

First, we produced a higher-coverage VCF. This VCF originally contained 616,477 markers, but was filtered to require that retained loci have data for 40\% or more of individuals and a minor allele frequency of 0.01. Markers were thinned such that only one marker on each contig was kept, so that all markers being analyzed were completely unlinked. This filtered VCF file contained 3052 markers, had a mean read depth of 33 reads per marker, and a median read depth of 20 reads per marker.

However, the filter initially applied for missing data may have excluded markers that were associated with sex. A sex-associated marker may only be present in about 40\% or less of the sample population, as the marker may not be present in one sex or every individual of that sex due to stochasticity in library preparation and sequencing. After re-filtering to adjust the missing data proportion to 0.2 (markers present in only 20\% or fewer individuals were excluded) the new VCF file contained 16020 markers, had a mean depth of 11 reads per marker, and a median depth of 8 reads per marker. 

We also aligned raw data to a draft \textit{de novo} genome assembly for the northern redbelly dace (\textit{Chrosomus eos}; Schultz \& Mandeville, unpublished data). Although this reference genome is more distantly related to creek chub and we therefore receive a lower alignment rate than aligning to the artificial creek chub reference genome (95\% alignment to the artificial reference genome and 73\% to the dace genome), aligning to a reference with larger scaffolds might be preferable for this analysis, as it could allow better identification of physically linked loci associated with sex determination \citep{Gopalakrishnan2017TheGenomics}. We performed the alignment and created a VCF file with the same methodology as above, with a missing data allowance of 40\%. This VCF file originally contained 1,233,215 markers, which dropped to 1132 after filtering, and had a mean read depth of 25 and a median read depth of 15. 

\subsection{Investigation of homomorphic sex chromosomes and polygenic sex determination}

As there is some evidence that RADSex might work less well in datasets with lower average sequencing depth \citep[as detailed in the methods paper describing RADSex;][]{Feron2021a}, we also applied a number of other analyses to identify potential sex-specific or sex-associated loci. We next performed a GWAS using GEMMA, a single-locus GWAS software \citep{Zhou2012}. Essentially, GEMMA treats each marker as a polymorphic allele and searches for two genotypes present exclusively in either sex. Therefore, GWAS would be able to identify markers from homomorphic sex chromosomes or regions from polygenic sex determination, but not heteromorphic sex chromosomes.

GEMMA requires genotype data at all markers, therefore, imputation of missing genotypes must be done beforehand. This increases statistical power compared to eliminating markers without genotypes for all individuals, which would drastically reduce the total number of markers used in the analysis \citep{Guan2008PracticalMapping}. We used the software entropy for imputation of missing genotypes, as it estimates posterior genotype probabilities at each locus, for each individual \citep{Gompert2014b, Shastry2021Model-basedMixed-ploidy}. The phenotype file contains a numeric character for the trait of each individual \citep{Zhou2012}. As the trait being examined, sex, is binary, 0 represented male and 1 represented female. The annotation file contained the information for each marker: the marker ID, base position, and contig number, sorted in the same order as the genotype file \citep{Zhou2012}. A relatedness matrix is first created that shows a measure of how related each of the individuals are to one another, then this file is used to run the univariate linear mixed model association test \citep{Zhou2012}. 

GEMMA results were visualized in RStudio, using a custom R script to create a Manhattan plot (Figure 2). The threshold for this plot is set at -log10(5e-6). Loci with p-values  above this threshold on the Manhattan plot are considered associated with sex at a statistically significant level and could indicate regions of the genome that are part of either a homomorphic sex chromosome or a  polygenic sex determination system.


\subsection{Analysis of genetic differentiation}

Regions of the genome associated with either a homomorphic sex chromosome or polygenic sex determination are expected to be highly differentiated between males and females of a species relative to other regions of the genome. To complement GWAS analyses, we calculated Weir and Cockerham locus-specific F$_{ST}$ using VCFtools \citep{Danecek2011TheVCFtools, Weir1984EstimatingStructure}. 

We used Rosner’s test to identify significant outliers from the Weir and Cockerham F$_{ST}$ values  \citep{DanDan2013a}. While the test is innately two-tailed and the outliers being examined only exist in the upper tail, the test is still suitable as any negative values returned will be excluded because they are not biologically relevant. As well, the data set fits the test’s assumption of a normal distribution. The test was performed for all three data sets in RStudio, using the {\tt rosnerTest} command from the package, {\tt EnvStats}. The value of alpha (p-value) was set at the default 0.05. The output is described in the relevant section of Results. 


\subsection{Discriminant analysis of principal components}

We completed an additional analysis, a discriminant analysis of principal components (DAPC) \citep{Jombart2010DiscriminantPopulations}, through the R package {\tt adegenet} \citep{Jombart2008Adegenet:Markers} to identify loci that contribute strongly to genetic differences between male and female individuals \citep[similar to][]{Junker2020a}. We first identified sex-associated loci using DAPC, then examined patterns of heterozygosity at the loci most strongly associated with differentiation between sexes.  


\subsection{Linkage disequilibrium of F$_{ST}$ and DAPC outlier loci}

Linkage disequilibrium was quantified by calculating the squared coefficient of correlation ($r^2$) between pairwise comparisons of all markers using VCFtools \citep{Danecek2011TheVCFtools, Pritchard2001LinkageData}. Using RStudio, we then took subsets of the data that only contained markers deemed outliers from DAPC and the Rosner's Test on F$_{ST}$ values. A Whitney-Mann \textit{U} test was performed on the data, using the function {\tt wilcox.test}, to assess whether the means of the subsets were significantly different from those of the entire relevant dataset, therefore undergoing less recombination than the rest of the genome \citep{Mann1947, Wilcoxon1945}. This was done for all 3 VCF files, resulting in 6 calculations that are summarized in Table\ \ref{tab:ld_means}. 

%-----------------------------------------------------------------------------------


\section{Results}

%-----------------------------------------------------------------------------------

\subsection{Using RADSex and GWAS, no markers were found to be associated with sex}


Using Radsex and GEMMA we found no loci significantly associated with sex (Fig.\ \ref{fig:radsex} and \ \ref{fig:manhattan}). Additionally, for RADSex, there were no markers with alleles shared by more than 19 of the 39 females or 21 of the 41 males. 


\subsection{Weir and Cockerham F$_{ST}$ analysis revealed some locus-specific genetic differentiation between sexes}


The 99th quantiles for each the 1132-, 3052-, and 16020-marker data sets were 0.136, 0.079 and 0.112 respectively, while the average  F$_{ST}$ values were all approximately 0 (Fig.\ \ref{fig:fst}). A Rosner’s test performed on the 16,020-marker data set showed that the markers with the largest 97 F$_{ST}$ values, ranging in value from 0.130 to 0.379, were statistically outliers that deviated from a normal distribution of F$_{ST}$, with p-value $<$ 0.05. 
Similarily, for the 3052- and 1132-marker F$_{ST}$ data sets, the 20 highest F$_{ST}$ values, ranging in value from 0.087 to 0.224, and the 13 highest F$_{ST}$ values, ranging in value from 0.134 to 0.300, respectively, were found to be statistical outliers which deviated from a normal distribution of F$_{ST}$, p-value $<$ 0.05.


\subsection{Values from F$_{ST}$ and GWAS had a weak, negative relationship}


A scatter plot was created for all three data sets to examine potential correlation of F$_{ST}$ and GWAS p-values. Ideally, some of the markers with F$_{ST}$ values determined to be outliers may have correlated positively with the markers with higher scoring, albeit non-significant, values from GEMMA. This was not the case, as the values were weakly correlated, often in the negative direction, when a simple linear regression was performed using R (Fig.\ \ref{fig:gwas_fst}). The 1132-, 3052-, and 16,020-marker data sets had a regression line with a slope of 0.298, -0.246, and -0.193, a p-value of 0.335, 0.496, and 0.036, and 1127, 2008, and 15961 degrees of freedom, respectively. While one of these findings is statistically significant, they are all unlikely to be biologically significant. 


\subsection{Discriminant analysis of principal components}

DAPC analyses identified a small number of loci that are differentiated between sexes in each data set. For the alignment to the artificial reference genome, 128 loci were significantly associated with sex in the 16,020 locus data set, and 27 loci were significantly associated with sex in the 3,052 locus data set. For the data set aligned to the dace reference genome (1132 markers), only 5 loci were significantly associated with sex. These loci were spread across 5 different scaffolds, rather than co-localized on a single large scaffold, and are therefore not consistent with placement on a differentiated sex chromosome or in proximity to a master sex determining gene. 


\subsection{Linkage disequilibrium was slightly higher among F$_{ST}$ and DAPC outlier loci}

In each data set, a small number of overlapping loci were identified as both Fst outliers and outliers in the DAPC analyses (Fig.\ \ref{fig:venn}). The subsets of outlier markers from F$_{ST}$ analysis had similar read depth to the average for the entire set: 11, 44, and 27 read depths on average for the subsets compared to 11, 33, and 25 for the whole 16,020-, 3052-, and 1132-marker data sets, respectively. Whereas the subsets of DAPC outliers had much higher mean read depths of 41, 119, and 35, respectively.

Linkage disequilibrium among loci identified as sex-associated by F$_{ST}$ and DAPC analyses was slightly higher than for the entire data set on average, suggesting that these loci are more closely linked and may be physically closer together in the genome or otherwise inherited together. The $r^2$ values for all data sets follow a right-skewed, exponential distribution. For the original 16,020-marker data set, the mean $r^2$ value for the F$_{ST}$ outliers was 0.103 and the mean of the entire data set was 0.054. The Whitney-Mann \textit{U} test resulted in p-value = 0.003. The mean $r^2$ value for the DAPC outliers was 0.068 and the p-value from the Whitney-Mann \textit{U} test was $<$ 2.2e-16 (Table \ \ref{tab:ld_means}). Similarly, for the 3052-marker data set, the mean $r^2$ of the F$_{ST}$ subset is 0.206, the mean of the entire data set is 0.031 and the Whitney-Mann \textit{U} test resulted in p-value $<$ 2.2e-16. The mean $r^2$ value for the DAPC outliers was 0.035 and the p-value from the Whitney-Mann \textit{U} test was 3.745e-13 (Table \ \ref{tab:ld_means}). Therefore, while the means were not large, they were still all significantly higher for both data sets aligned to our artificial reference genome. For the 1132-marker data set, the mean overall $r^2$ value is 0.0449, the mean $r^2$ for significant DAPC values is 0.063, and the mean $r^2$ for F$_{ST}$ outlier values is 0.102. The difference was significant for the F$_{ST}$ outliers (p-value = 1.21e-12), but not for the outliers from DAPC (p-value = 0.711) (Table \ \ref{tab:ld_means}). Again, the LD values are quite low, but taken all together, LD  tended to be higher for the significant F$_{ST}$ and DAPC markers when compared to other markers in each data set.


%-----------------------------------------------------------------------------------

\section{Discussion}

%-----------------------------------------------------------------------------------
The study of sex determination presents significant challenges, because many clades, including some vertebrates, feature a diversity of sex determination systems \citep{Bachtrog2014a}. While mammals and birds follow a straightforward heteromorphic sex chromosome system, other vertebrate clades may use much more subtle means that remain elusive to our research methods \citep{Charlesworth1996TheCompensation}. Fish are particularly diverse in their systems of sex determination, and such systems are believed to have rapid evolutionary turnover in these lineages \citep{Heule2014, Pennell2018a}. 

In this context, it is not entirely unexpected that analyses of potential sex determining regions in this study produced equivocal results. Overall, while no statistically significant sex-associated markers were found in the analyses with RADSex and GEMMA, some loci identified by analyses of Fst outliers and discriminant analysis of principal components may be candidates for involvement in sex determination in creek chub. Based on these results, and because most North American cyprinid fish with known sex determination have some form of genetic sex determination, we believe that it is still likely that there is a genetic basis for sex determination in creek chub \citep{Ashman2014a}. Based on our analysis, it is likely that sex determination in creek chub may take the form of homomorphic sex chromosomes with a small sex determining region, or polygenic sex determination. 


\subsection{The sex determination region may be small or minimally differentiated}


Based on initial results, it is unlikely that creek chub possesses highly differentiated, heteromorphic sex chromosomes similar to an XX/XY or ZW/ZZ sex determination system. An area of the genome as large and differentiated as a heteromorphic sex chromosome would have likely been detected by at least a small number of markers using RADsex (Fig.\ \ref{fig:radsex}), had greater genetic differentiation between sexes (Fig.\ \ref{fig:fst}), or exhibited greater loadings or involved more loci in the DAPC analyses. 

The creek chub genome has 25 chromosomes, so on average, our data set would have contained over 100 markers per chromosome in the 3052-marker data set and over 600 markers per chromosome in the 16,020-marker data set \citep{Gold1987}. While this is still only a small fraction of the genome, as is typical of reduced-representation sequencing strategies like genotyping by sequencing and RADseq, it is unlikely that a chromosome as differentiated between the sexes as a heteromorphic sex chromosome would have gone completely undetected. 

Creek chub may instead have homomorphic sex chromosomes; however, the sex determining region would likely be quite small or scarcely differentiated, as it was not detected by GWAS and only a few loci were sex-associated in Fst and DAPC analyses. The same sentiment exists for polygenic sex determination, where there may be multiple, small sex determination regions across two or more chromosomes that evaded targeting by the sequenced markers. The latter is quite probable, as the loci of interest were scattered throughout the genome rather than localizing to one scaffold in the dace genome alignment. 

Additionally, these few, scattered sex-associated loci may suggest that sex determination systems are not a reproductive barrier for cyprinid fish. Sex chromosomes can change in fashion quickly and species with heteromorphic sex chromosomes are faster at reaching reproductive isolation than those with other forms \citep{Pennell2018a}. Haldane's rule also states in cases where hybrid individuals are inviable or infertile, it is more likely to be the heterogametic sex that is negatively affected \citep{Haldane1922}. Given that we find no evidence for heterogamy in creek chub, and as interspecific hybridization and production of fertile offspring is known to occur among cyprinid fishes including creek chub and a suite of closely related species, this suggests that their sex determination system is not a barrier to reproduction and that Haldane's rule is unlikely to be involved in the maintenance of reproductive isolation between cyprinid minnows \citep{Broughton2011Current, Dowling1993, Ross1981}. This further supports our evidence which suggests creek chub likely have small or minimally differentiated homomorphic sex chromosomes or a polygenic sex determination system.  


\subsection{Assessing the likelihood of environmental sex determination in creek chub}


Creek chub could also determine sex using external environmental influences, or a combination of environmental and genetic sex determination \citep[as in sea bass][]{Palaiokostas2015}. While most species with a known sex determination mechanism are exclusively subject to GSD, a fraction of reptile and fish species may undergo either ESD or GSD with effects of temperature on sex ratios (GSD + TE) \citep{Bachtrog2014a, Charnov1977, Muralidhar2018, Shen2019a}. In this process, external factors, such as temperature, pH, photoperiod, or social factors, induce epigenetic modification or hormonal targeting of sex determination genes \citep{Heule2014, Shen2019a}. This reduces gene expression, promoting development of an alternate phenotypic sex \citep{Heule2014, Shen2019a}. In the case of environmental sex determination in creek chub, there would be little to no sex-related genomic differences between males and females; therefore, all high LD, F$_{ST}$, and DAPC outliers values identified would just be coincidental. 

However, to the best of our current knowledge, ESD is almost non-existent in the Cyprinidae family and seen in only 15\% of bony fishes \citep{Ashman2014a, Bachtrog2014a}. While turnover of various GSD mechanisms is quite common within fish families \citep[for example in sticklebacks][]{Natri2019a}, an entire switch to ESD is not \citep{Heule2014, Pan2019, Pennell2018a}. Thus, while the possibility exists for creek chub to undergo ESD, particularly GSD + TE or a mixed system, it is most likely still the case that GSD is the mechanism of sex determination in these fish \citep{Muralidhar2018, Shen2019a}, but that either the genetic architecture of sex determination is too complex to conclusively identify it, or our dataset did not contain the relevant genomic regions. 

This sort of ambiguity about environmental vs. genetic sex determination is not unique to creek chub. Sex determination in lampreys, for example, has remained a mystery despite over 20 years of study. The mechanism has been suggested to be environmental, as there has been no large sex chromosomes found and there is some evidence for density-dependent sex determination \citep{Docker1994, McCauley2015LampreysEra}. Thus, while studies with equivocal results about the basis of sex determination may seem less satisfying than those that confidently identify a clear genetic basis of sex determination, studying more challenging species provides a unique opportunity to learn more about non-standard sex determination systems.  

\subsection{Technical limitations associated with the identification of sex determining regions}

Thus far, we have focused on biological interpretations of our results, but technical limitations may also contribute to the inability to detect the genetic basis of sex determination in some studies, including this one. Specifically, reduced-representation library preparation and sequencing methods like GBS, ddRAD, and RAD sequence only a small fraction of the genome, despite identifying thousands of single nucleotide polymorphisms (SNPs), so in any study like this, the loci of interest may be excluded \citep{Lowry2017BreakingAdaptation}. This becomes particularly relevant when attempting to identify the genetic basis traits that are controlled by many loci of small effect, like a polygenic sex determination system might be. 

As well, the power of this study may have been somewhat limited by our sample size of 79 individuals -- 39 males and 41 females. In a study on quantitative trait loci, which work in similar fashion to polygenic sex determination loci, power analyses on GWAS done by \citealp{Kardos2016Whole-genomePopulations} show that it may be difficult to identify loci that each work in small effect, without thousands of individuals. However, that is not to say that this sample size would be ineffective at identifying loci with large effect sizes, especially for a binary trait like sex in our sampled fish. For the binary trait of survival, \citealp{Margres2018Large-effectCancer} used a similar number of individuals for GEMMA -- 41 males and 69 females -- but were able to find 5 SNPs in females and 37 in males that explained 76\% and 43\% of phenotypic variance, respectively. Therefore, while our sample size still may not have been strong enough to detect multiple small polygenic regions, it would have likely wielded enough power to detect a large sex determining region or chromosome. 

Alternatively, our data set may have contained sex-associated loci that we were unable to detect statistically due to insufficient sequencing depth. Marker depth was extremely variable across loci, and many marker depths were relatively low, particularly in the 16,020-marker data set (Fig. S2 B). High marker depth was associated with results from DAPC, but not from Weir and Cockerham F$_{ST}$, suggesting that the results from DAPC may be somewhat biased towards markers with better coverage. While DAPC might be affected by read depth, F$_{ST}$ doesn't seem to be, so those markers that stick out from both DAPC and F$_{ST}$ analyses might be more reliable than just the ones from DAPC.

Additionally, there were no markers in the RADSex output tile plot that were shared by all individuals, likely because the read depth was low and variable enough that not all markers had even one read for all individuals (Fig.\ \ref{fig:radsex}). This was not the case for the any of the 7 species with shown tile plots in the methods paper describing the software \citep{Feron2021a}, therefore, RADSex may not have been able to accurately assess the creek chub sequencing data, leading to a null result. Issues with sequence depth at individual loci may have arisen with GEMMA as well. However, methods like DAPC have been successfully used in to identify sex-associated loci in similar data sets \citep[e.g. in Lake Tanganyika sardines][]{Junker2020a}, so it is difficult to parse how much data quantity/quality contributes to this result relative to actual complexity of sex determination systems. It is clear that methods for the detection of sex determination systems are still in need of development, especially for studies of non-model organisms without reference genomes and lower-coverage data sets that prioritize sampling more individuals over sequencing at higher coverage, as is the case for many conservation-associated studies.  


\subsection{Future directions}


While GSD is the most likely method of sex determination in creek chub, to date, no studies have been done on this species to rule out the effects of temperature or other environmental influences on sex ratios of creek chub offspring. ESD is unlikely to be the only mechanism behind sex determination in this species, however, temperature effects may play a role \citep{Muralidhar2018, Shen2019a}. Additionally, development of a creek chub genome assembly would allow for genome-wide linkage mapping and assessment of gene ontology in areas of recombination suppression, as done by \citealp{Kamiya2012b}, as MSD genes are often transcription factors involved in development and maintenance of reproductive organs \citep{Herpin2015, Pan2019}. Further research into sex determination in creek chub and other minnows in the \textit{Cyprinidae} family will likely yield insight into the evolutionary diversification and current dynamics of interspecific hybridization in this species-rich group of fishes. 


%-----------------------------------------------------------------------------------


\bigskip

\noindent \textbf{\large Data accessibility} 

Upon the acceptance of this manuscript, data and scripts used for analysis will be made publicly available in Data Dryad and the NCBI Sequence Read Archive. These materials will be made available to reviewers upon request. 

%-----------------------------------------------------------------------------------

\pagebreak


\bibliography{references.bib}
\bibliographystyle{mol-ecol.bst}


\clearpage

%-----------------------------------------------------------------------------------

\section{Figures}

%-----------------------------------------------------------------------------------


\begin{figure}
    \centering
    \includegraphics[scale=0.85, angle=0]{figures/radsex_output.png}
    \caption{Tile plot depicting the output from RADSex. The number of females with a given marker are displayed on the y-axis and number of males with a given marker on the x-axis, with number of markers held by that number of males and females indicated by darkening colour. Tiles containing markers that are found in only one sex at a statistically significant level would be shown surrounded by a red box, however, no such markers were found. }
    \label{fig:radsex} 
\end{figure}

\clearpage

%-----------------------------------------------------------------------------------


\begin{figure}
    \centering
    \includegraphics[scale=0.6, angle=0]{figures/manhattan_vertical.png}
    \caption{Manhattan plots displaying output from Wald Chi-Squared Test implemented in GEMMA. The -log10(p-value) for each marker’s association with sex is displayed on the y-axis. The significance threshold is represented by a solid line, at -log10(5e-6). No significant markers were found in any of the three data sets. A: 3052-marker data set. B: 16,020-marker data set. C: 1132-marker data set with chromosome numbers along the bottom for the 25 largest contigs. Chromosomes 1 to 4 and 7 are not shown as no markers mapped to these regions. Chromosome 0 contains markers from all small, excess contigs that did not map to any other chromosome. }
    \label{fig:manhattan} 
\end{figure}


\clearpage

%-----------------------------------------------------------------------------------


\begin{figure}
    \centering
    \includegraphics[scale=0.45, angle=0]{figures/fst_all_vertical.png}
    \caption{Weir and Cockerham FST values. Solid line indicates 0 and dashed line indicates 99th quantile. A: Histogram of the 3052-marker data set. B: Scatter plot of the 3052-marker data set. C: Histogram of the 16,020-marker data set. D: Scatter plot of the 16,020-marker data set. E: Histogram of the 1132-marker data set. F: Scatter plot of the 1132-marker data set. }
    \label{fig:fst} 
\end{figure}


\clearpage
%-----------------------------------------------------------------------------------


\begin{figure}
    \centering
    \includegraphics[scale=0.45, angle=0]{figures/venn_all.png}
    \caption{Statistically significant outliers from Fst analysis and DAPC. Markers that overlap between the Fst outliers and DAPC outliers have been identified as loci of interest for future analyses. A: 3052-marker data set. B: 16,020-marker data set.  C: 1132-marker data set.}
    \label{fig:venn} 
\end{figure}


\clearpage

%-----------------------------------------------------------------------------------

\begin{table}
\caption{Linkage disequilibrium expressed as mean $r^2$ values and one standard deviation for the 3 data sets and subsets of outlier markers from Fst and DAPC. Values that are statistically different from the mean of the entire data set is denoted with a star. }
\vspace{0.25in}
\begin{tabular}{p{1.2in}|p{1.5in}|p{1.5in}|p{1.5in}}

  {\bf Data set} & {\bf Overall mean} & {\bf Fst outlier loci} & {\bf
                                                                  DAPC outlier loci
                                                                  }\\
\hline
  3052-marker & 0.0313  $\pm$ 0.0891 & 0.2064 * $\pm$ 0.1671 & 
                                                        0.0350 * $\pm$ 0.0487 \\
& & \\
16020-marker & 0.0541 $\pm$ 0.1373 & 0.1030 * $\pm$ 0.1497 & 0.0678 * $\pm$ 0.1373 \\
& & \\
1132-marker & 0.0449 $\pm$ 0.0821 & 0.1021 * $\pm$ 0.1055 & 0.0629 $\pm$ 0.0842 \\

\label{tab:ld_means}
\end{tabular}
\end{table}


\clearpage


%-----------------------------------------------------------------------------------


\end{linenumbers}
\end{document}
